# AFM-RL: Large Protein Complex Docking Using AlphaFold-Multimer and Reinforcement Learning

**DOI:** 10.1101/2024.01.20.576386

**Authors:** Tunde Aderinwale, Rashidedin Jahandideh, Zicong Zhang, Bowen Zhao, Yi Xiong, Daisuke Kihara

## Abstract

Various biological processes in living cells are carried out by protein complexes, whose interactions can span across multiple protein structures. To understand the molecular mechanisms of such processes, it is crucial to know the quaternary structures of these complexes. Although the structures of many protein complexes have been determined through biophysical experiments, there are still many important complex structures that are yet to be determined, particularly for large complexes with multiple chains. To supplement experimental structure determination, many computational protein docking methods have been developed, but most are limited to two chains, and few are designed for three chains or more. We have previously developed a method, RL-MLZerD, for multiple protein docking, which was applied to complexes with three to five chains. Here, we expand the ability of this method to predict the structures of large protein complexes with six to twenty chains. We use AlphaFold-Multimer (AFM) to predict pairwise models and then assemble them using our reinforcement learning framework. Our new method, AFM-RL, can predict a diverse set of pairwise models, which aids the RL assembly steps for large protein complexes. Additionally, AFM-RL demonstrates improved modeling performance when compared to existing methods for large protein complex docking.

## Introduction

Protein complex interactions play a crucial role in various cellular processes. These interactions can range from a simple pair of proteins to a complex network involving multiple proteins. The availability of the three-dimensional structure of these protein complexes is essential for understanding their interactions and functions. While experimental methods exist for determining the structure of protein complexes, it becomes increasingly difficult as the size of the complex grows. Not only is it costly and time-consuming, but it can also be technically challenging to experimentally determine the structure of large protein complexes. This is reflected in the limited number of experimentally solved structures of large protein complexes in the proteomes of humans and other organisms.

Over the years, computational methods have been used to predict the protein structures of complexes of various sizes. Pairwise protein docking, which involves a protein complex with two chains, has been studied extensively ^1–6^, with impressive results shown in the Critical Assessment of PRedictions of Interactions (CAPRI)^7^ Competitions. Methods for multimeric protein docking ^8–10^, which involves complexes with more than two chains, have also been developed, but they have limitations such as restrictions on the size of the protein complex, usually between 3 to 6 chains, and limitations in the type of symmetry supported ^11–14^. These methods also often require extra information to simplify the problem.

The recent breakthrough of AlphaFold ^15^ for single protein structure prediction has given rise to numerous methods ^16–19^ extending it to the prediction of multiple protein complexes. One such example is ColabFold ^16^, where input sequences were concatenated with linker sequences. Another example is AF2Complex^20^, which predicts the interaction between multimeric protein complexes using the original AlphaFold method without retraining or paired MSA. Subsequently, Deepmind released AlphaFold-Multimer^21^, an adaptation of the original framework trained to predict the structure of multiple protein docking. The method was applied to protein complexes of two to three chains. we also developed a multiple-chain docking method, Reinforcement Learning for Multimeric protein docking with LZerD ^22^ (RL-MLZerD). RL-MLZerD was applied to protein complexes with three to five chains.

There is still a very limited number of computational methods for predicting the structure of large protein complexes. Complexes with six or more chains are still very challenging to predict due to the combinatorial explosion when such protein interactions are considered. AlphaFold-Multimer can be directly applied to predict the structure of large protein complexes, however, as shown in Bryant et al. ^23^, there are two main challenges with this approach. First, the accuracy decreases as the number of chains increases. The second challenge is the GPU memory limitation, as a protein complex of 2500 sequences would quickly fill 40GB of memory. These two challenges highlight the need for a more robust approach to predicting the structure of large protein complexes. MoLPC ^23^ is an example of such an approach. They showed that one can predict the structure of the protein complex by first decomposing it into smaller subcomponents, and then assembling the full structure by searching and combining the subcomponents. As highlighted in their results, their approach quickly runs into two problems. The first is missing interfaces within the subcomponents, where the critical interface that is needed to accurately predict the large protein complex is missing within the subcomponents predicted. The second is high clash and overlaps of protein chains between subcomponents.

In this work, we introduce AFM-RL (Fig. 1), which similarly approaches the problem as our previous work, RL-MLZerD. Large protein complexes are assembled by first using AFM to predict the structure of pairwise docking poses (decoys). Next, the decoys are selected and assembled. The key addition of this work is the forced sampling induced into AlphaFold-Multimer to ensure critical interfaces are not lost when the large complex is decomposed into small pieces while preserving high-quality pairwise prediction. Our results showed that AlphaFold-Multimer is forced to predict alternative interfaces and diversify the potential candidates when presented with multiple copies of the same protein chain. AFM-RL was benchmarked on a dataset of 57 protein complexes with six to twenty (6-20) chains. The average root mean square deviation (RMSD) and TM-SCORE of the best-assembled model across all targets are 10.09 Å and 0.81, respectively. AFM-RL showed a better performance when compared to MoLPC with an average RMSD of 10.09 Å compared to 18.63 Å of MoLPC.

**Fig. 1.**
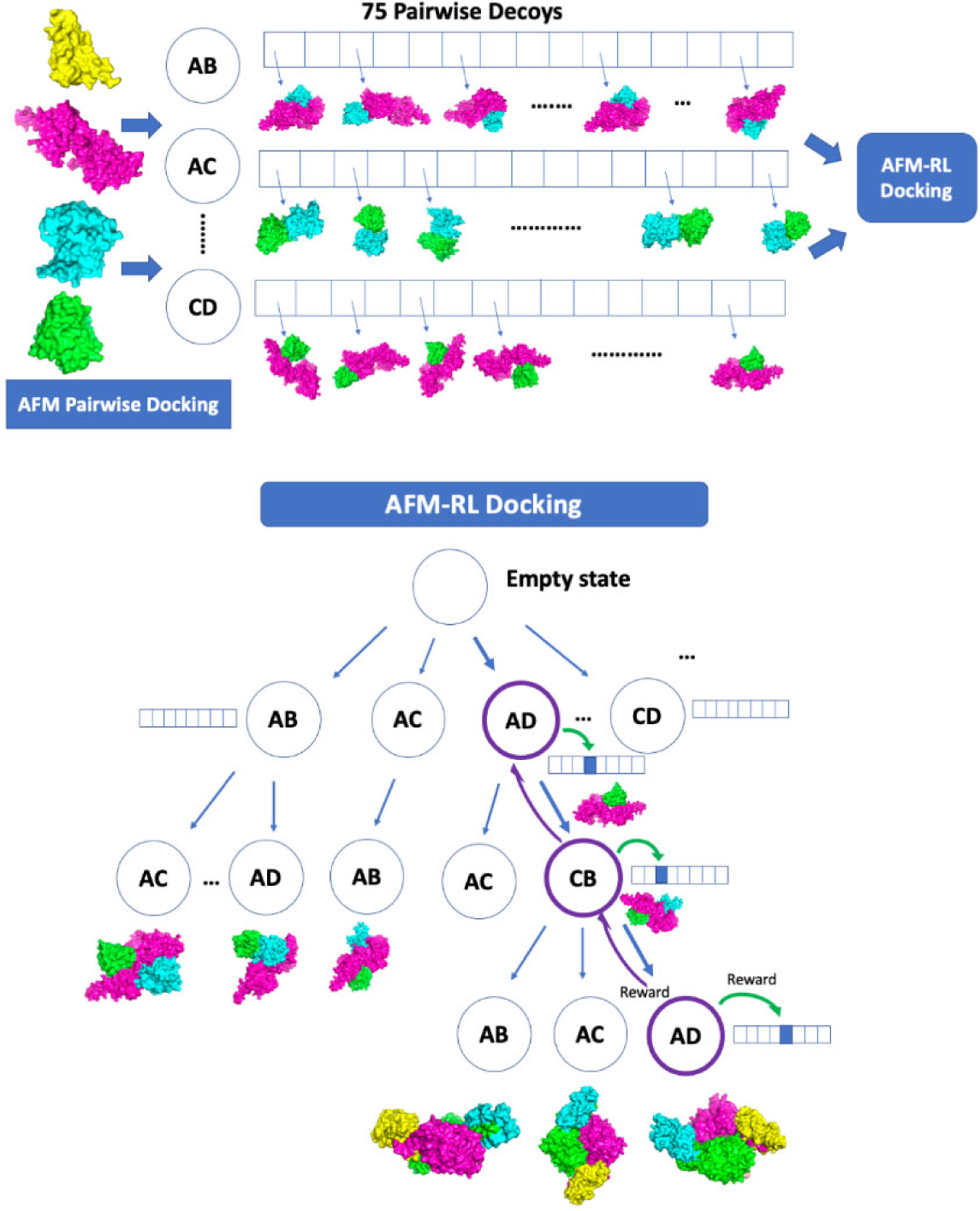
Overview of AFM-RL. In the first step, pairwise docking of every pair of subunits is performed using AlphaFold-Multimer (AFM). 75 decoys are selected for each pair. In the subsequent step, pairwise decoys are assembled using reinforcement learning. When an assembly of a full subunit complex is successful without too many clashes, the reward may be propagated back to assembly states and decoy states used along the episode.

## Results

### Pairwise docking quality impact

The success of any assembly method depends on the quality of the pairwise models. If there are no good quality pairwise models, the assembly stage of the process would be searching within the pool of bad models. Fig 2a shows the effect of the quality model on the final assembled complex. The y-axis is the RMSD of the resulting complex that was assembled using the pairwise models. The x-axis is the interacting pairwise coverage percentage. To determine the interacting pairwise coverage percentage, we select the pairs of chains that are interacting (any pair within a 5 Å radius compared to the native structure). For these interacting pairs, we count how many have a minimum of one predicted model within 4 Å of the same pair in the native structure. We then report the percentage of interacting pairs that meet these criteria. For example, if we have 4 interacting pairs and 3 of them have a predicted model that is within 4 Å of the native structure, the x-axis for that example would be 75%. We can observe from Fig 2a that as the coverage percentage increase, the resulting complex model quality also increases (lower RMSD). There is one outlier example in the plot, 1VF7 with interaction coverage percentages of 57.90% and RMSD of 43.61 Å. Despite capturing 57.90% of the interacting pair, the RMSD of the assembled complex is 43.61 Å. The complex is a 13 chains homomer which two layers of 7 and 6 chains, both forming a long cyclical structure. The predicted structure assembled the first layer as 9-interacting chains and the remaining 4 chains were placed around the top of the first layer, resulting in the high RMSD.

**Fig. 2.**
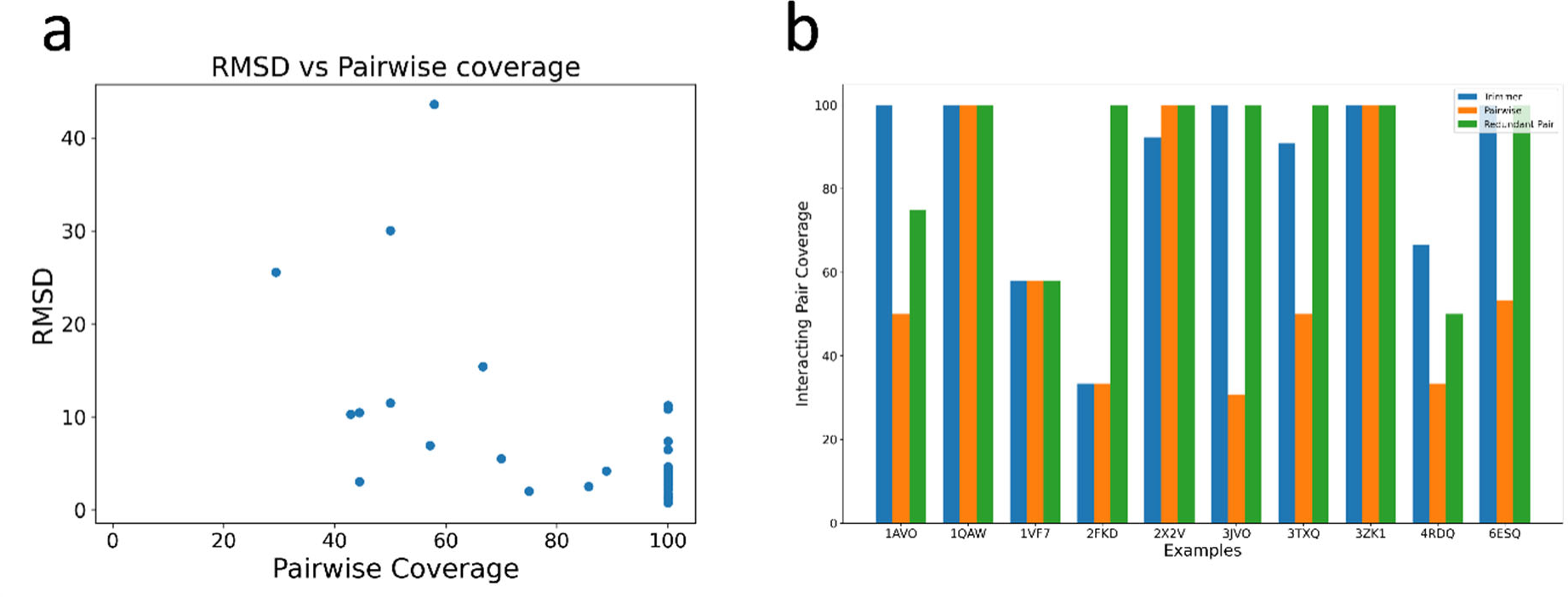
Pairwise model impact on complex assembly. **a.** Comparison of complex RMSD against the interacting pairwise coverage. The y-axis is the complex RMSD, the x-axis is the pairwise coverage percentage for each target in the dataset. **b.** Comparison of forced sampling approach compared to regular pairwise and trimmer approach.

Next, we evaluate the performance of the forced sampling (redundant pair) approach compared to the regular pairwise and trimeric approaches. In the forced sampling approach, we use AFM to predict the subcomponents for all combinations of pairs. For each pair, we present AFM with three copies of the second chain. For example, for A and B chains, we provide AFM with A1B3. We then extract all the different sets of A1B1 predicted interactions. The regular pairwise approach predicts one copy for both chains, i.e A1B1. Lastly, the trimeric approach involves selecting three chains as subcomponents from all possible combinations and then extracting pairwise models from the predicted trimeric model. Fig 2b shows the interacting pairwise coverage percentage of the three approaches. For this comparison, we randomly selected 10 examples. The redundant pair approach captured 100% of the interactions for 7 of the examples. Trimmer approach captured 100% interaction for 5 examples and lastly the regular pairwise approach captured 100% of the interaction for 3 examples. The trimeric approach had two example where it captured more of the pairwise interactions, 4RDQ and 1AVO (66.67%, 100.00%) compared to the redundant approach (50.00%, 75%). Overall, the average interacting pairwise coverage for the redundant pair, regular pairwise, and trimeric approaches is 88.29%, 60.87%, and 84.11%, respectively.

### Comparison with AlphaFold-Multimer

Out of the 57 complexes in the dataset, we were able to predict 42 with AFM, but the remaining 15 could not be predicted end-to-end on a 40GB GPU workstation due to an Out of Memory error, highlighting the limitations of full end-to-end prediction. On average, AFM predicted structures with an RMSD of 5.53 Å and a TM-score of 0.91, compared to 5.45 Å and 0.87 for AFM-RL. When comparing individual cases, AFM predicted structures with a lower RMSD for 27 examples, compared to 15 for AFM-RL. Individual results are shown in Supplementary Table S1 and S2.

Overall, end-to-end prediction with AFM is effective when the structure is small enough to fit into memory and the conformation is simple. However, for large and complex structures with numerous chains that cannot be predicted end-to-end, AFM-RL is able to assemble the structure accurately, emphasizing the importance of gradually assembling the complex from its subcomponents.

### Comparison with MoLPC

First, we explain the comparison criteria for this section. Full complex RMSD is calculated by using all of the CA atoms in the complex without discarding any of them. Partial RMSD, on the other hand, is using MMAlign to calculate the RMSD and TM-score. In this case, we refer to the RMSD as partial RMSD because some CA atoms from the structure are discarded. This usually occurs when the best alignment found by MMAlign corresponds to the core part of the complex.

We show the full complex RMSD (Fig 3, panel A) of AFM-RL compared to MoLPC for all the examples in our dataset. On average, the best-assembled complex of AFM-RL has an RMSD of 10.09 Å compared to 18.63 Å of MoLPC. When we compare individual cases, AFM-RL has 50 examples with lower RMSD compared to MoLPC. There are 5 cases (excluding 2BX9) where MoLPC has a lower RMSD compared to AFM-RL. Additionally, there are 3 examples (3TXQ, 2BX9, and 1TYF) where AFM-RL can assemble the complex to completion, but MoLPC is only able to assemble the structure partially: 3TXQ: 10 out of 11 chains, 2BX9: 3 out of 14 chains, and 1TYF: 7 out of 14 chains.

**Fig. 3.**
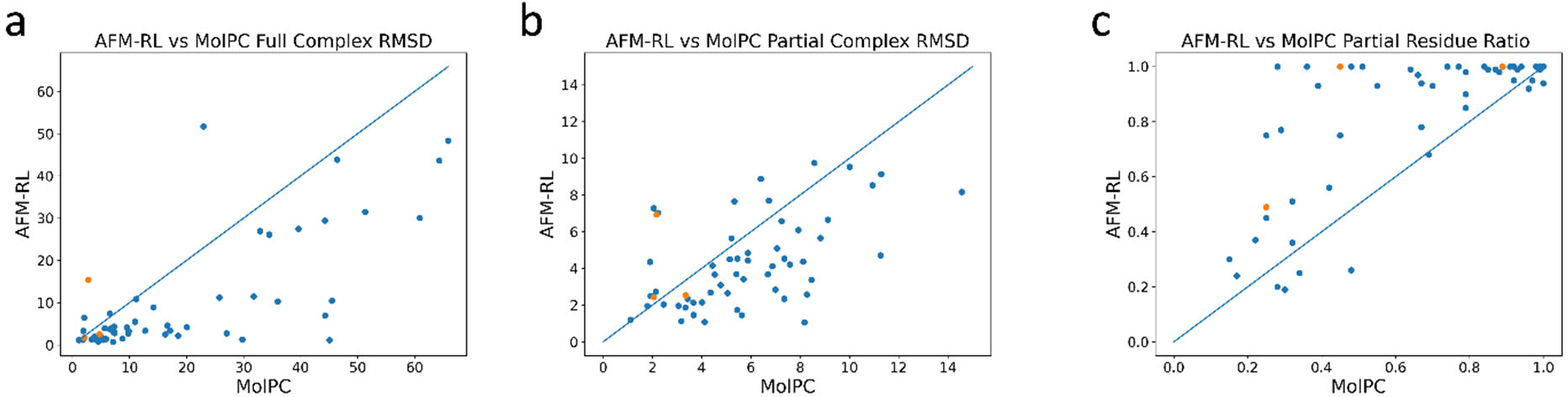
Comparison of AFM-RL results against MoLPC. **a**, The full RMSD of the predicted complex. **b**, The partial RMSD of the predicted complex. **c**, The ratio of residue covered in the partial RMSD calculation. For all the panels, the AFM-RL result is on the y-axis, and MoLPC is on the x-axis.

Next, in Fig 3b, we compare the partial RMSD of AFM-RL against MoLPC. On average, AFM-RL has a 4.26 Å RMSD compared to 5.71 Å of MoLPC with 0.81 and 0.69 TM-score, respectively. Similarly, there are 43 individual examples where AFM-RL has a lower RMSD compared to 12 (excluding 2BX9 and 3TXQ) of MoLPC. Finally, when we evaluate the ratio of the sequence included in the partial RMSD and TM-score calculation, on average, AFM-RL has a ratio of 81% compared to 66% of MoLPC, highlighting the overall quality of the modeled structure from AFM-RL. This is because MMalign was able to superimpose more residues from the AFM-RL model compared to MoLPC.

Furthermore, we rank and score all the models assembled by AFM-RL and compare the top 5 models against the best-assembled model from MoLPC (we compare against the best model because MoLPC only produces a single output model). On average, the top 5 models of AFM-RL have a 16.48 Å RMSD compared to 18.63 Å of MoLPC for the full complex RMSD. The average partial RMSD of AFM-RL is 3.96 Å compared to 5.71 Å of MoLPC, with a 0.78 TM-score compared to 0.69 of MoLPC. Finally, a partial residue ratio of 82% compared to 66% of MoLPC (Fig 3c).

### Example of docking models

Figure 4 shows four examples of structures assembled by AFM-RL. The models are shown with each chain in a different color, while the native structure is in gray. 1MGQ (Fig 4a) and 6LYP (Fig 4c) are both 7-chain complexes. All 7 chains interact cyclically. Overall, AFM-RL assembled both structures with an RMSD of 1.34 Å and 2.79 Å, respectively. Figure 4, panel B is 7AB3, a 6-chain complex. The overall structure forms a U-shape with two chains interacting on the left, right, and center. Both AFM and AFM-RL were able to assemble the structure with RMSD of 1.05 Å and 2.16 Å, respectively. However, MoLPC predicted a cyclical structure. The interaction interface of 2 pairs was predicted incorrectly and created additional interaction between another pair, resulting in an overall complex RMSD of 18.51 Å. The last example is 1M3U (Fig 4d), a 10-chain complex, with a Dihedral symmetry. The structure breaks into two layers of 5 chains with each layer forming cyclical interactions. AFM-RL assembled the structure with the lowest RMSD of 1.2 Å, followed by AFM with 1.3 Å, and finally MoLPC with an RMSD of 5.44 Å.

**Fig. 4.**
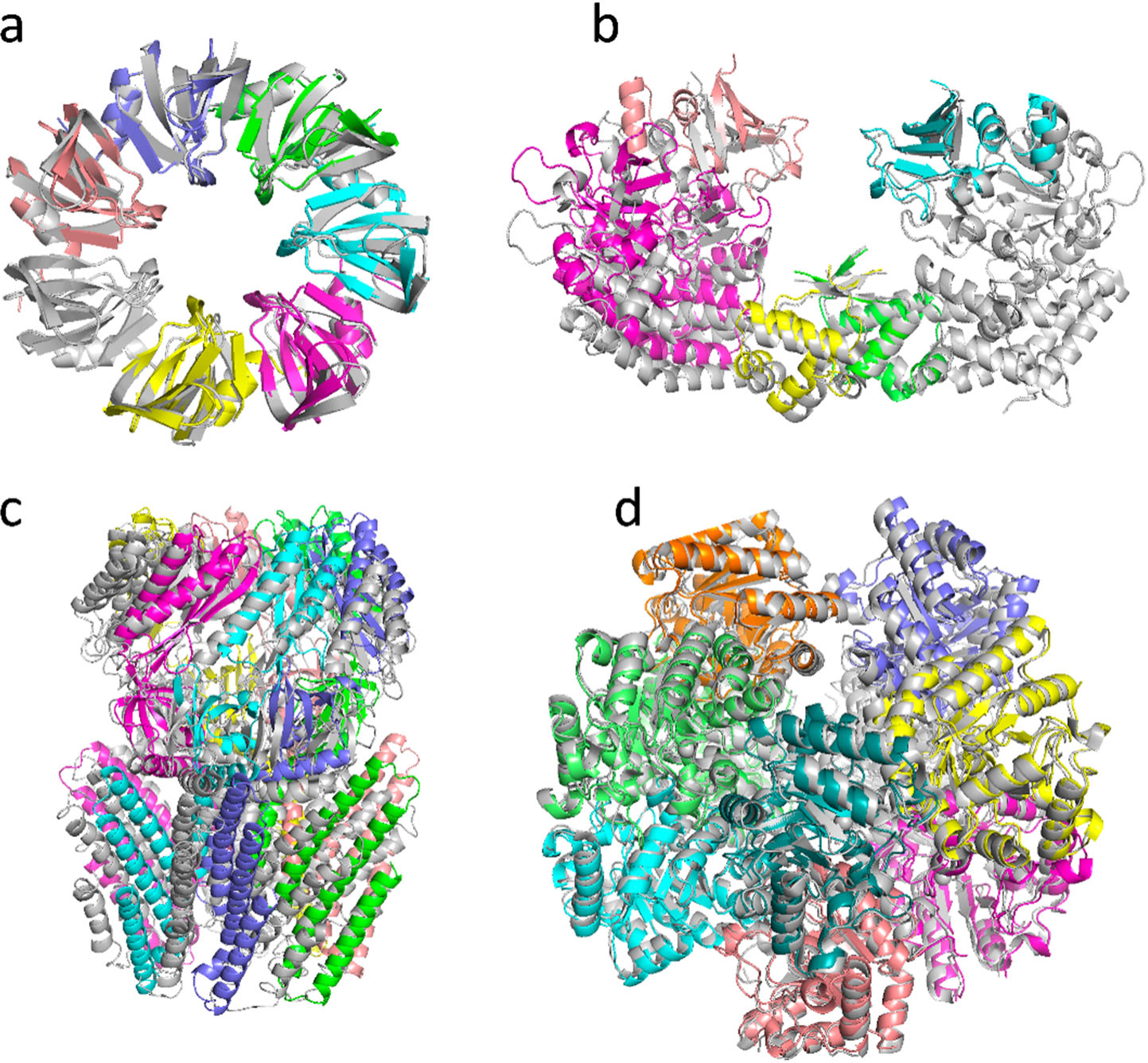
Examples of AFM-RL predicted models. The native structure solved by an experimental method is in grey, and the generated models are shown in colors. Each chain is colored in a different color. **a**, Structure of a heptameric sm-like protein from methanobacterium thermoautotrophicum, PDB ID: 1MGQ. A 7-chain complex. RMSD, 1.34 Å. **b**, Structure of E. coli toxin-antitoxin system HipBST (HipT S57A), PDB ID: 7AB3. A 6-chain complex. RMSD, 2.16 Å. **c**, Structure of mechano-sensitive ion channel protein 1, mitochondrial wild type, PDB ID: 6LYP. A 7-chain complex. RMSD, 2.79 Å. **d**, the Crystal structure of E. coli ketopantoate hydroxymethyl transferase complex, PDB ID: 1M3U. A 10-chain complex. RMSD, 1.20 Å.

Next, Fig 5 highlights two examples of a complex with non-typical conformations and how each method assembled the complex. The first example is 4ZWS (Fig 5 a-d), a 7-chain complex. It is an alpha-helix-rich complex with C1 symmetry. However, chains F and G (gray and purple color) form an opening with a rotation of around 120°. Fig 5a shows the native structure, Fig 5b shows the AFM prediction, Fig 5c shows the AFM-RL assembled structure, and Fig 5d shows the MoLPC structure. AFM predicted a cyclical structure and did not capture the opening of the two chains. AFM-RL perfectly captured the opening and rotation of the two chains. MoLPC predicted a non-cyclical structure and failed to capture the rotation of the two chains. Overall, AFM-RL assembled the lowest RMSD structure with 2.51 Å RMSD compared to 9.81 Å and 16.25 Å of AFM and MoLPC. The second example is 7NAK (Fig 5 e-h), an 8-chain complex with H symmetry. Fig 5e shows the native, Fig 5f shows the predicted AFM, Fig 5g shows the AFM-RL assembled structure, and Fig 5h shows MoLPC. Both AFM and AFM-RL were able to predict the complex structure with RMSD of 1.15 Å and 1.14 Å, respectively. However, MoLPC predicted a cyclical structure, resulting in an RMSD of 45.05 Å.

**Fig. 5.**
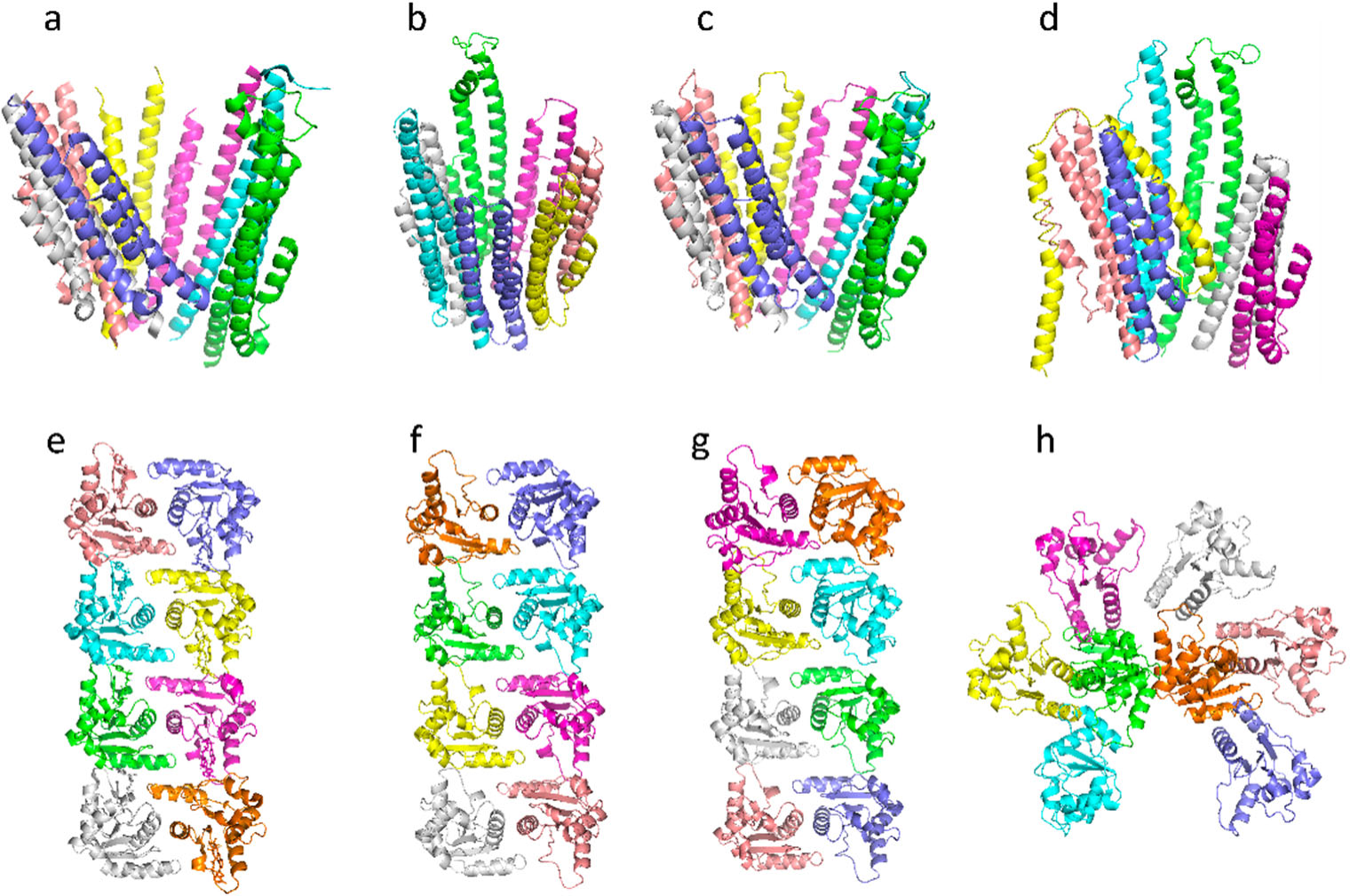
Example case study of a predicted model from all methods. The first example (panel a-d) is the Crystal structure of the bacteriophage T4 recombination mediator protein UvsY, PDB ID: 4ZWS. A 7-chain complex. **a.** The native structure was solved by experiment. **b.** The predicted structure by AFM, RMSD 9.81 Å. **c**. Predicted structure by AFM-RL, RMSD 2.51 Å. **d.** The predicted structure by MoLPC, RMSD 16.25 Å. The second example (panel e-h) is the structure of activated human SARM1, PDB ID: 7NAK. An 8-chain complex. **e.** The native structure was solved by experiment. **f.** Predicted structure by AFM, RMSD 1.15 Å. **g.** Predicted structure by AFM-RL, RMSD 1.14 Å. h. Predicted structure by MoLPC, RMSD 45.05 Å.

Finally, Fig 6 shows two examples where AFM-RL can assemble the complex completely without facing several clashes of the chains or missing critical interaction interfaces. The first example is 1TYF (Fig 6, a-c), a 14-chain complex. It forms 2 layers of 7 chains each, the top layer interacting with the bottom layer through the beta-strand in the middle of each chain in the layer. The interaction is small and is the only connection between the layers. Fig 6a shows the native complex, Fig 6b and Fig 6c show the predicted structure by AFM-RL and MoLPC, respectively. AFM-RL successfully assembled the full complex (RMSD 2.53 Å) as our forced sampling approach captured the critical interaction between the layers. However, MoLPC was only able to assemble the bottom half of the layer (Fig 6c) with the 7 chains having RMSD of 4.79 Å. The next example is 3TXQ (Fig 6, d-f), an 11-chain complex. All the chains interact cyclically. Fig 6d shows the native, Fig 6e and Fig 6f show the predicted structure by AFM-RL and MoLPC, respectively. AFM-RL was able to predict the full complex structure without one chain clashing with another. However, MoLPC was able to assemble 10 chains and couldn’t fit the last chain into the complex without it clashing with another chain in the complex. Overall, AFM-RL’s full 11-chain complex has an RMSD of 1.71 Å compared to 2.11 Å of the 10-chain MoLPC predicted complex.

**Fig. 6.**
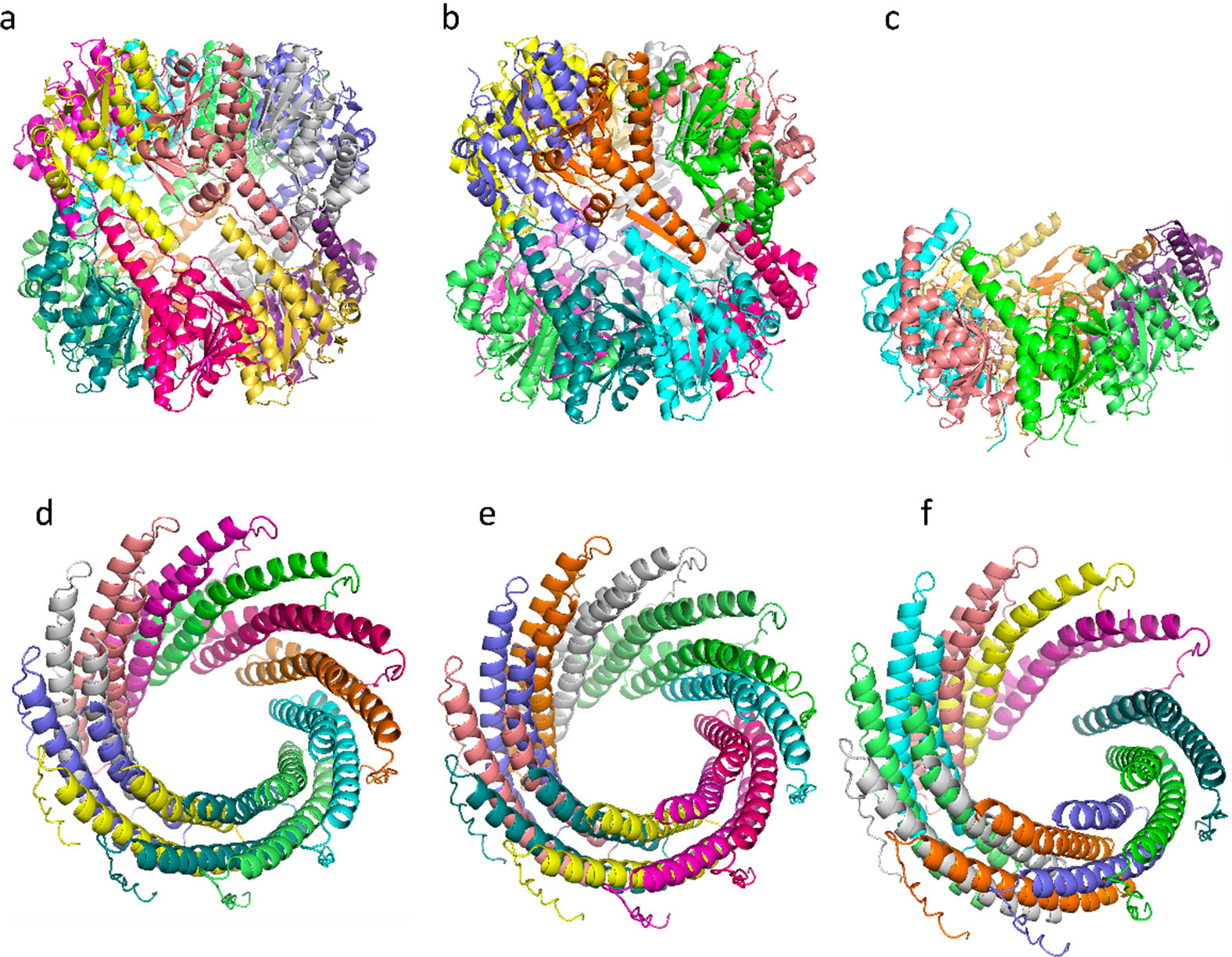
Example case study of completed complex prediction by AFM-RL compared to the partial prediction by MoLPC. The first example (panel a-c) is the structure of ATP-dependent Clp protease proteolytic, PDB ID: 1TYF. A 14-chain complex. **a.** The native structure was solved by experiment. **b.** The Predicted structure by AFM-RL, RMSD 2.51 Å. **c.** The predicted structure by MoLPC, RMSD 4.79 Å for 7 chains out of 14. The second example (panel d-f) is the structure of phage 44RR small terminase gp16, PDB ID: 3TXQ. An 11-chain complex. **d.** The native structure was solved by experiment. **e.** Predicted structure by AFM-RL, RMSD 1.71 Å. **f.** Predicted structure by MoLPC, RMSD 2.11 Å for 10 out of 11 chains.

## Discussion

Regular AFM was not trained to predict the structure of large complexes; However, it can still be used for the prediction of large complexes with good quality. The main limitation of this end-to-end prediction approach is the intensive GPU memory requirement. Even with a 40 GPU memory, some examples from this dataset could not fit into memory. While this is a major limitation for regular AFM, our method, AFM-RL, does not have this limitation. AFM-RL can be used to predict the structure of large protein complexes by decomposing the structure into a small set of sub-structures and then assembling them step-by-step, which avoids the GPU memory limitation.

In this work, we show how AFM can be extended with RL to predict the structure of large complexes. Using our forced sampling approach, AFM generated a diverse set of pairwise structures that resulted in multiple interfaces for a single pair of proteins. The average interacting pairwise coverage percentage for the examples in our dataset is 84%. On average, our predicted complex structure has a lower RMSD of 10.09 Å and a higher TM-score of 0.81 compared to MoLPC’s RMSD of 18.63 Å and a TM-score of 0.69 across all examples.

The predicted structure from our method can be used to aid experimental structural predictions. Experimentalists can first run our method to gain insight into the full complex structure and its subcomponents, plausible complex conformations, pairwise interactions, and topology. All of these findings can be used to reduce the complexity of their experiments.

The future direction of this work would be to apply the method to large protein complexes across multiple organisms. This large-scale application would generate thousands of needed 3D structures of protein complexes, which will aid in understanding biomolecular activities and interactions in cells. Finally, we plan to develop a self-service portal where users can submit the sequence of their protein complex and our servers run the prediction pipeline, generating downloadable models from an interactive web browser.

## Acknowledgments

This work was partly supported by the National Institutes of Health (R01GM133840, R01GM123055, and 3R01GM133840-02S1) and the National Science Foundation (CMMI1825941, MCB1925643, DBI2146026, and DBI2003635). The National Science Foundation of China (grants nos. 61832019 and 62172274). The author would like to thank Purdue Rosen Center for Advanced Computing (RCAC) in West Lafayette, Indiana for providing computational resources. We also acknowledge the support from the Purdue Institute of Inflammation, Immunology, and Infectious Disease (PI4D). The computational experiments were partially run at the Center for High-Performance Computing, Shanghai Jiao Tong University.

## Author contributions

DK conceived the study. TA designed the algorithm, developed the codes, performed the benchmark studies, and associated analyses. RJ was involved in performing some parts of the benchmark studies, ran AFM and MOL-PC and provided example cases. BZ, ZZ ran AFM for the complex structure and the trimmer studies. TA wrote the initial draft of the manuscript and DK edited it. All authors have read and approved the manuscript.

## Methods

### Dataset construction

We evaluated AFM-RL on the same dataset as MoLPC. We selected complexes with 6 to 15 chains. Due to the computation intensity requirement and limited GPU availability, we limited the number of selected examples from each chain to a minimum of three and randomly selected the 3 example complexes. For 15 chains, there are only 2 examples; we show results for only one example out of the two. AFM resulted in an Out-Of-Memory (OOM) error for the second example during the pairwise step. We have 57 complexes in total. The PDB list for all the complexes is provided in Supplementary Table S1.

### Pairwise docking

AFM-RL builds protein complex models in two steps. First, it constructs a pool of pairwise decoys for each pair of chains. In the second step, it searches through the pool of pairwise decoys and explores different combinations as assembly episodes within the RL framework. The resulting assembled models are evaluated, and rewards are propagated in the RL’s Q-table, which records preferred decoys with high probability scores, to guide efficient searches for assembly.

In the pairwise docking stage, AlphaFold-Multimer (AFM) was used to generate candidate decoys for each pair of chains. For each pair, we ran AFM with duplicated copies of one of the chains in order to force AFM to expand sampling and provide more than one interface for the pair. For example, instead of presenting AFM with an input fasta file of A1B1, we provided A1B3. This ensures that AFM provides more than one candidate interface and interaction site between A and B. This forced sampling is used to diversify the candidates for each pair produced by AFM. Finally, we ran AFM with the replication option, which outputs 25 candidate pairs, which are then multiplied by 3 from the duplicated copies, resulting in 75 candidates for each of the pairs in the complex.

### Assembling pairwise decoys with RL

Similar to our previous work, the RL framework shown in Fig 1. is used to assemble the generated pairwise decoys. An RL episode is denoted as a docking process, the pseudocode of which is provided in Fig. 7. which makes choices of two types of states along the path. The first one is called assembly states (circles in the diagram in Fig. 1), which denote subunit combinations, e.g. AD > AC (adding C to the complex by choosing a decoy of the AC pair). Starting from an empty state at the top of the diagram, subunits are added one at a time by choosing a pairwise decoy until a terminal state is reached, where all subunits are assembled. The second type of state is called decoy states (an array of boxes in Fig. 1), which denotes decoys in the pool generated for each subunit pair. Thus, an episode consists of a set of selections of assembly states and a decoy state at each assembly state.

**Fig. 7.**
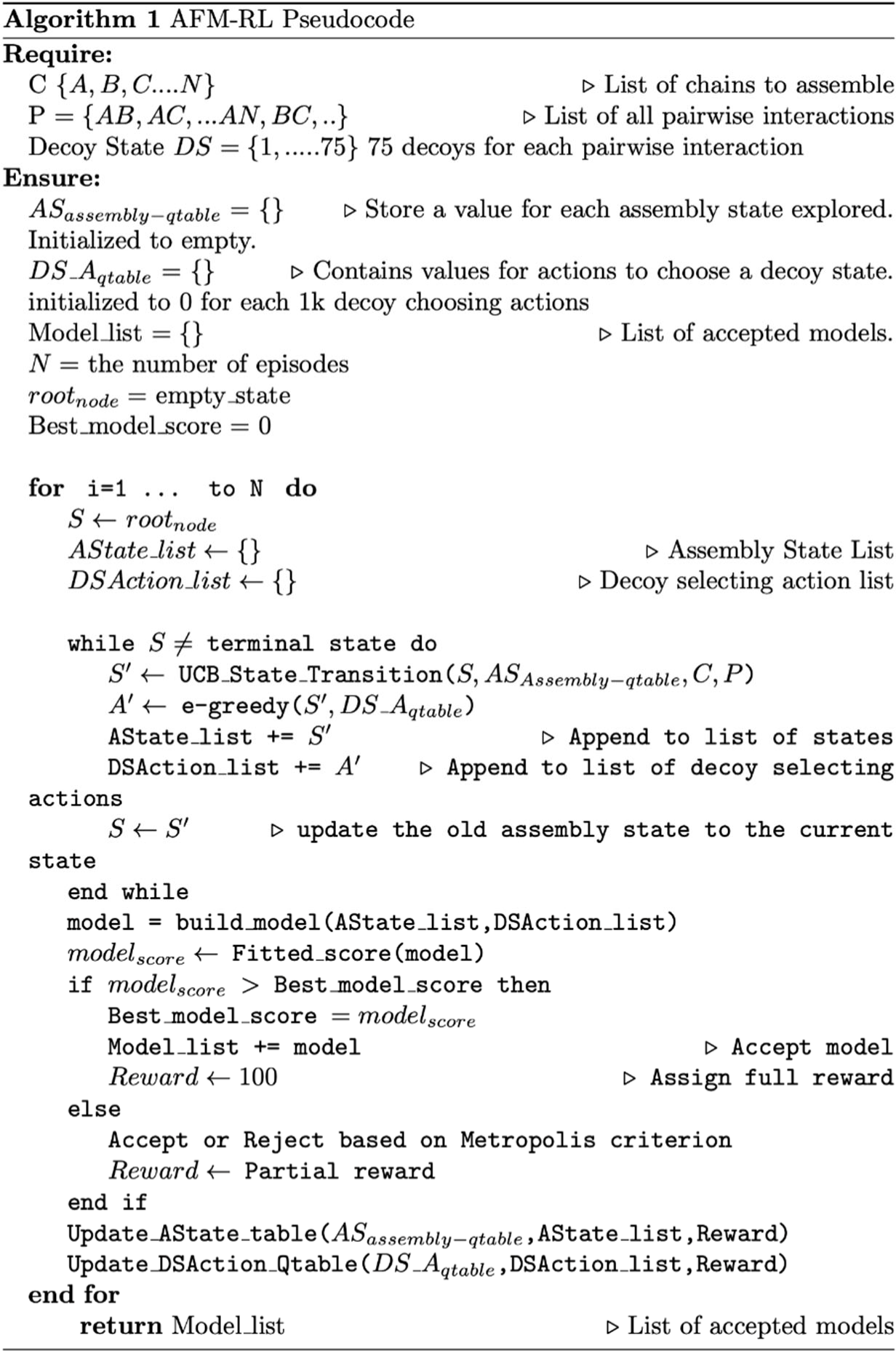
Pseudocode of AFM-RL. For each episode, *AState_list* represents the assemble state visited for the episode, *DSAction_list* represents the selected actions (decoys) for each decoy state. The *build_model* function takes the *AState_list* and *DSAction_list* and returns the assembled model. *Fitted_score* computes a score of a model. *Update_state_table* updates the state value estimate for the assemble state that participated in building the model, *update_action_qtable* uses the Bellman Fold equation to update the decoy state actions value estimate that participated in the model building.

Accordingly, there are two types of actions, one that selects the next assembly state and the other that specifies a decoy from a decoy pool for a subunit pair. The next assembly state is selected with the Upper Confidence Bound (UCB) policy ^24^, which tries to balance exploitation and exploration with the following score:

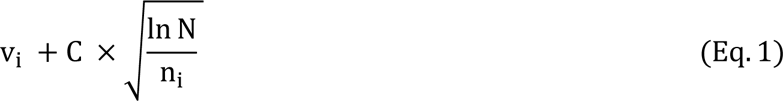

where v_i_ is the value of assemble state i, C is a hyper-parameter, which was set to 1.0 at the beginning and slowly reduced by 1.0/(the total number of episodes) after each episode is performed. N is the total number of visits to the parent state of the assemble state i, n_i_ is the number of visits to the assembly state i. When N is small at the early stage of computation, states are selected mainly by v_i_, while less-visited states are more explored as N becomes larger. At the beginning of the computation when values are not accumulated yet, the next assembly state is randomly chosen.

The second type of action deals with the decoy state. Each time an assembly state is visited, a decoy-state is selected from a pool using e-greedy ^25^ as the policy. The e-greedy policy dictates that the agent exploits the best action (i.e. the decoy with the highest shape score) among possible choices most of the time (i.e. 1-e) but randomly selects a decoy-state with probability e. We set e to 1.0 at the beginning and slowly reduced it by 1.0/(the total number of episodes) after each episode was performed.

Here we briefly walk through the algorithm (Fig. 1, 7). The docking procedure starts from a root node in Fig. 1, and the next assembly state is selected according to the UCB policy, followed by a selection of a decoy-state. This procedure is iterated until the full complex is built. The developed full chain complex model is evaluated by a scoring function that is a linear combination of four scoring terms, a molecular mechanics potential ^9^, the solvent accessible surface area, the radius of gyration (RG), and atom clash counts. RG is included to encourage compact assemblies. An atom clash is recognized if two heavy atoms are closer than 3 Å to each other. Weights of the terms were determined by a logistic regression classifier with 4-fold cross-validation trained on complex models of two quality classes, with an interface RMSD (as defined in the CAPRI evaluation ^7^) less than or over 4.0 Å.

During an episode, a partially-assembled complex is checked for atom clashes due to the newly selected action at the decoy state. If the resulting partially-assembled complex at the state has several atomic clashes higher than the threshold (n-1) * 100, where n is the number of assembled protein chains, the newly selected decoy state is rejected and replaced with a different decoy that is ranked among the top 5 by Q score until an acceptable model is found. The modeling process moves on to the next assembly state to add the next subunit if the top 5 decoy selections still result in a model with a high number of atomic clashes.

Once an episode generated a complex model, the model evaluation score is checked to see if it has the best score that has been discovered thus far from all episodes. This signifies a newly discovered model with the best score, and a such model is assigned a full reward of 100 points. Any other model score that falls short of this criterion is assigned a partial reward if the Metropolis criterion is met:

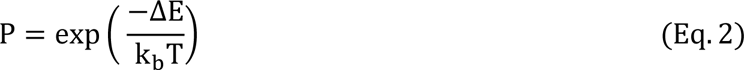

Where ΔE is the score difference between the new model and the current best model. We report the results with a normalization factor, k_b_T, which was set to 6.0. A complex model is accepted if P is larger than a random number generated between 0 and 1. As a variation of the method, we also report results where we set P to a constant value of 0.6, thereby giving a constant 60% chance of acceptance for assembled model regardless of the model score.

If a model score is close to the current best, it has a high chance to be accepted even if it is lower than the best. If the model is accepted, a partial reward is assigned by discounting the 100 points based on the calculated probability to reflect the difference between the current best score and the score of the new model. On the other hand, a reward of 10 points is provided if the model does not pass the metropolis criterion. This is because the model is geometrically possible (if it is not rejected by a high number of atom clashes) and thus we do not want to penalize the path that generated the model. A penalty reward of -2 is assigned if the model is rejected due to a high number of atomic clashes. Thus, unlike conventional RL methods where the goal is fixed, the goal of AFM-RL is constantly being updated based on the current best-discovered model.

Parameters used in RL, namely, the normalization factor (Eq. 2) set to 6.0, the full reward set to 100, the reward of 10 given to a rejected model, and the penalty of -2, were determined by a small number of tests on a couple of targets. We showed in our previous work how the different parameters affect the results.

At last, we explain how values are updated in the RL framework. Values for assembly states, v_i_ used in Eq. (1), in eligible states are updated at the end of an episode as follows:

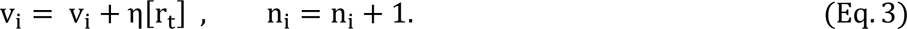

Where η is the learning rate, set to 1.0; r_t_ is the reward assigned to the complex model at the end of the episode (e.g. 100 points). n_i_is the number of times the assembly state was visited. The eligible states that are updated are those which participated in the model building path of the episode. According to Eq. (3), the same reward value is added to all the eligible states along the episode.

The reward obtained at the end of the episode is also propagated to the decoy states selected. The update is based on the temporal difference using the Bellman-Ford Equation ^2^^6^:

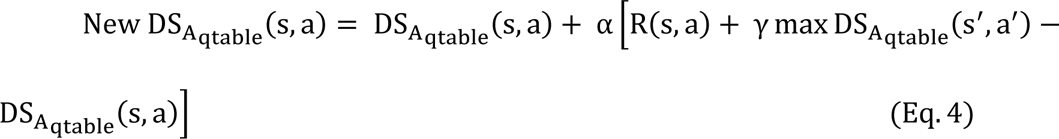

where s is the assembly state, and a is the selected action (decoy) for the assembly state. New DS_A_qtable_(s, a) is the updated Q score for the assembly-decoy state pair, DS_A_qtable_(s, a) is the old Q score, α is the learning rate, where we use an adaptive learning rate which decays from 1.0 to 1.0 * 0.85^(episode/1000)^ after every 1000 episodes. R(s, a) is the reward. For a terminal state, it is the reward given to the episode and it is 0 for all intermediate states. γ is the discount rate for future rewards, which was set to 0.25. max DS_A_qtable_(s′, a′) is the maximum score among decoys (a′) of the next state visited s’.

Typically, 1500 to 12,000 models are generated for a target complex. They are clustered by LB3DClust ^27^, and are then ranked by the sum of score ranks by the scoring function mentioned above and the VoroMQA score ^28^. For each cluster, the best scoring model was selected as the representative.

## Data availability

The dataset used in this work is also made available at https://doi.org/10.5281/zenodo.7623376.

## Code availability

The source code for AFM-RL is available at https://github.com/kiharalab/AFM-RL

