## Supplementary material for "AFM-RL: Large Protein Complex Docking Using AlphaFold-Multimer and Reinforcement Learning": Supplemenary file caption

**Supplementary file S1 (separate csv file). Benchmark dataset details metrics information**

This file contains the raw metric value for all 57 targets. Column A is the PDB ID of the target, Column B is the number of chains in the complex structure. Column C-D is the RMSD value for AlphaFold best and top 5 structure. Column E-F is the RMSD value for AFM-RL best and top 5 structure. Column G-H contains the partial RMSD value for AFM-RL best and top 5 structure respectively. Column I contain the RMSD value of MolPC predicted structure. Column J is the partial RMSD structure of MolPC predicted structure. Column K-L is the percentage of the sequence covered in the partial RMSD calculation for both MolPC and AFM-RL, finally column M contain the percentage of the sequence covered by the top 5 predicted structure of AFM-RL.

**Supplementary file S2 (separate csv file). Raw TM-SCORE result for all methods.**

The raw TM-score value for all 57 targets is listed here. The first column is the PDB ID of the target, column B is the number of chains in the complex structure. Column C-D is the tmscore of AlphaFold, AFM-RL and MolPC predicted complex structure respectively. Column E is the tmscore from the top 5 scoring structure of AFM-RL.
